# Alternative interaction sites in the influenza A virus nucleoprotein mediate viral escape from the importin-α7 mediated nuclear import pathway

**DOI:** 10.1101/447979

**Authors:** Patricia Resa-Infante, Jaume Bonet, Swantje Thiele, Malik Alawi, Baldo Oliva, Gülsah Gabriel

**Affiliations:** Heinrich Pette Institute, Leibniz Institute for Experimental Virology, Hamburg, Germany; Structural Bioinformatics Lab (GRIB), Universitat Pompeu Fabra, Barcelona Research Park of Biomedicine (PRBB), Barcelona, Spain; Bioinformatics Core, University Medical Center Hamburg-Eppendorf, Germany; Center for Structural and Cell Biology in Medicine, University of Lübeck, Germany

**Keywords:** Influenza virus, host-pathogen interaction, virus evolution, nuclear transport, protein-protein interaction

## Abstract

Influenza A viruses are able to adapt to restrictive conditions due to their high mutation rates. Here, we addressed the question by which mechanisms influenza A viruses may escape restriction by the cellular importin-α7 protein, a component of the nuclear import machinery required for avian-mammalian adaptation and replicative fitness in human cells. Therefore, we assessed viral evolution in mice lacking the importin-α7 gene. Here, we show that particularly three mutations occur with high frequency in the viral NP protein (G102R, M105K and D375N) in a specific structural area upon *in vivo* adaptation. Moreover, our findings suggest that the adaptive NP mutations mediate viral escape from importin-α7 requirement likely due to the utilization of alternative interaction sites in NP beyond the classical nuclear localization signal and importin-α isoforms. However, viral escape from importin-α7 is, at least in part, associated with reduced replicative fitness in human cells.

## INTRODUCTION

Influenza A virus progeny production involves various interactions with viral and cellular factors in the infected cell [1, 2]. Since viral transcription and replication takes place in the cellular nucleus, the viral polymerase complex needs to adapt to the nuclear import machinery of the new host upon interspecies transmission [3, 4].

Importin-α is a constituent of the classical mammalian nuclear import pathway. It acts as an adaptor protein that recognizes nuclear localization signals (NLS) of cargo proteins which are then transported together with the importin-β_1_ receptor as a ternary complex into the nucleus [5]. The nuclear import of incoming viral ribonucleoprotein complexes (vRNPs) is believed to occur mainly by the interaction of the viral nucleoprotein (NP) with importin-α isoforms [6, 7].

Two NLSs have been described for NP: one on the N-terminal region (NLS1: residues 1-13) and a bipartite NLS in the middle of its globular domain (NLS2: residues 198-216) [8-10]. Inhibition of either site significantly reduces the nuclear import of vRNPs, albeit NLS1 seems to play a major role. However, blocking both NLS still allows residual nuclear accumulation of NP, suggesting the presence of other NLS regions which additionally mediate nuclear import [11].

Mammalian cells contain several importin-α isoforms that are specialized for the nuclear import of NLS-containing cargo proteins [12]. Among those, mammalian influenza viruses specifically require importin-α1 and importin-α7 for efficient vRNP nuclear import and increased replicative fitness in human cells [13, 14]. Importin-α7 further contributes to the development of influenza virus mediated pneumonia and increased mortality in murine models [13-15]. Conversely, importin-α7-knockout (α7^−/−^) mice are resistant to 2009 pandemic H1N1 influenza A virus (2009 pH1N1) infection, while all wild-type (WT) mice present 100 % lethality.

We have previously described that influenza viruses are able to escape the restricting conditions in mice lacking the importin-α7 gene. There, we could identify mutations in the viral genome that mediated enhanced virulence in mice [16]. In this study, we investigated the mechanisms behind the accelerated *in vivo* viral evolution that leads to increased pathogenicity in mice.

## RESULTS

### Accelerated viral evolution in importin-α7^−/−^ mice reveals accumulation of mutations in the viral NP gene

Escape from usage of the nuclear import factor importin-α7 upon enforced influenza virus adaptation to α7^−/−^ mice by serial lung-to-lung passaging results in enhanced virulence by an alternative, yet unknown molecular pathway [16]. In order to elucidate the underlying mechanisms resulting in increased disease development, we have assessed viral evolution *in vivo*. Therefore, we have isolated total RNA from infected lungs from α7^−/−^ mice after passage 5 when 100 % lethality was observed (**Fig. 1A**) [16]. Deep sequencing revealed that the highest variance per position in the original virus was in the range of 0.1-1.0 % (**Fig. 1B**, black dots). However, some positions showed single nucleotide polymorphisms (SNP) with a variance higher than 10 % upon viral adaptation (**Fig. 1B**). The high-frequency SNPs detected in PA, HA and NA viral genes were described before and mediate enhanced viral replication and pathogenesis [16]. Besides, the high-frequency SNP at position 183 of PB1 is a synonymous mutation and one SNP was found at nucleotide position 632 of the NS gene that translated to the R211K mutation in the viral NS1 protein. This residue is located in the C-terminal tail of NS1 which possesses a high degree of sequence and length variation [17]. However, most mutations that occurred upon serial *in vivo* passaging accumulated in the viral NP gene and showed highest frequency along successive infections (**Fig. 2A**). Namely, NP G102R described before and the herein newly identified mutations NP M105K and NP D375N. Interestingly, NP_G102R_ and NP_M105K_ mutations, which are in close proximity in the sequence of NP, were not present in the same RNA molecule during sequencing, suggesting independent occurrence in viral quasispecies. All of these three NP host adaptive mutations are clustered at the surface of the NP body domain (**Fig. 2B**).

**Figure 1.**
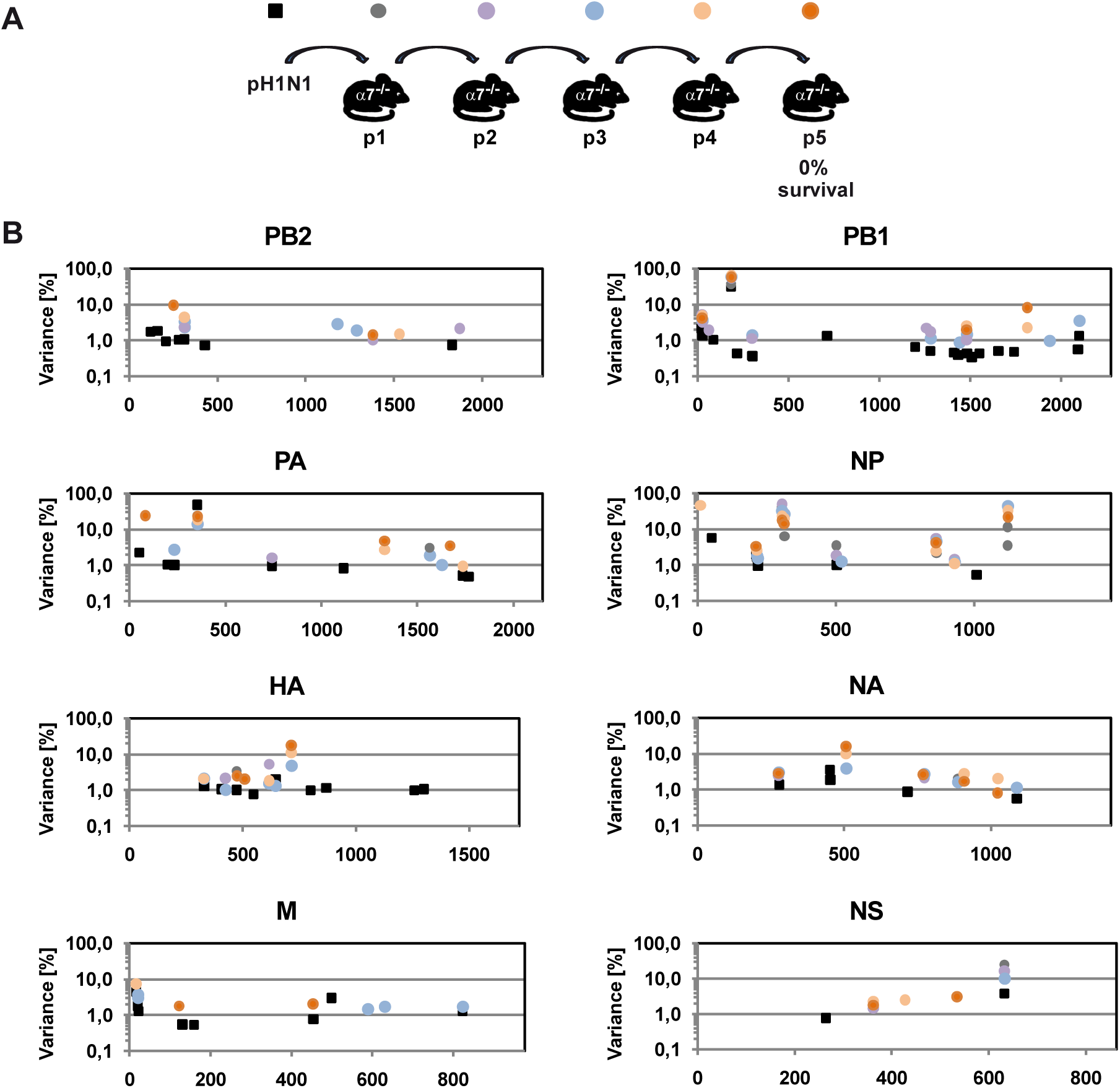
Deep sequencing reveals low sequence variation during adaptation of pandemic pH1N1 influenza A virus in a restrictive environment. **A**. Schematic experiment overview for sequential lung-to-lung passaging. Three α7^−/−^ mice were intranasally infected with 10^5^ p.f.u. of a pH1N1 strain (A/Hamburg/NY1580/09). Three days post infection, supernatants of the lung homogenates from these infected mice were pooled (named p1) and used to infect a second group of three α7^−/−^ mice. This procedure was repeated five times until 100 % lethality of the resultant virus was obtained. The lung homogenates from every passage were named p1, p2, p3, p4, and p5, respectively. **B**. Virus sequence variance of each passage was measured by deep sequencing. Individual RNA sequence reads were mapped to a consensus sequence derived from the isolate A/Hamburg/NY1580/09 (H1N1). The base pair positions relative to the reference sequence of each viral gene are indicated on the x-axis. SNPs were enumerated as described in Methods. No variants were called below 0.1 % of variance. The frequency of variants is presented as black squares for the parental strain (pH1N1) and as colored circles for the passages, specifically grey for p1, purple for p2, blue for p3, light orange for p4 and dark orange for p5.

**Figure 2.**
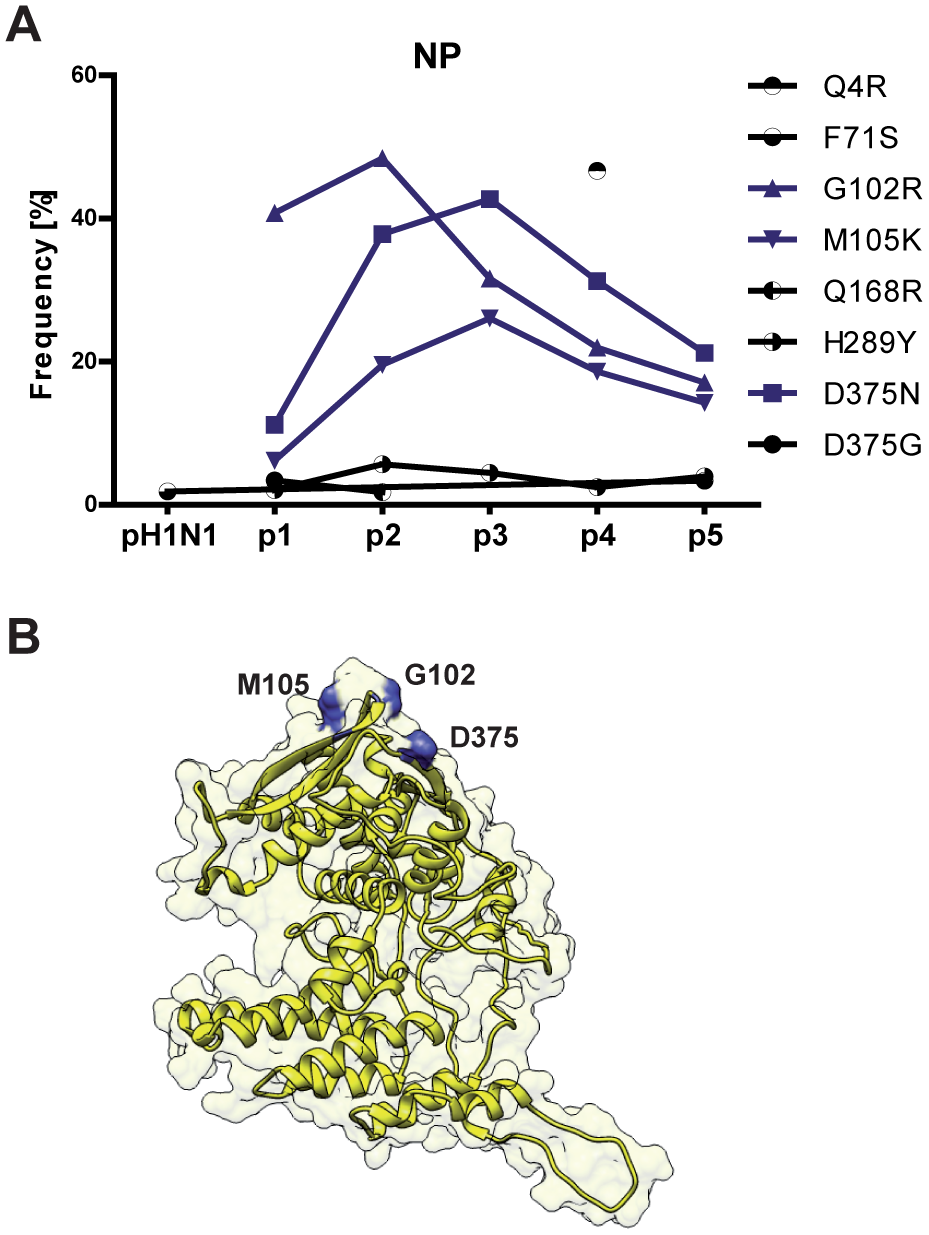
The viral NP gene accumulates mutations during adaptation of pH1N1 virus to α7^−/−^ mice. We focused our studies on NP, the genomic segment with a variance above 25 % that originated from non-synonymous substitutions. **A**. Frequency of the mutations appearing in viral NP during passaging. G102R, M105K and D375N mutations depicted in blue showed the highest frequency during passaging. **B**. Structural model of pH1N1 NP. The localization of the three mutations of interest is highlighted in blue.

These results highlight a specific structural area in NP that accumulates mutations during adaptation to importin-α7 host factor absence.

### Viral escape mutations in NP alter vRNP activity and virus replication

To study the effect of these NP host adaptive mutations on viral polymerase activity, we reconstituted viral RNPs in human epithelial cells (**Fig. 3A**). Although none of the adaptive NP mutations affected the biological activity of the recombinant RNPs, all of them are localized in a structural domain previously described to be involved in determining resistance of the viral RNP to the antiviral effect of human myxovirus resistance protein A (MxA) [18]. Thus, we tested the effect of MxA expression on the activity of reconstituted viral RNPs (**Fig. 3B**). RNP activity in the presence of antivirally inactive MxA-T103A was used to normalize the data obtained with functional MxA. Here, we observed that the NP_G102R_ and NP_M105K_ mutations led to reduced vRNP activity in the presence of MxA, in contrast to the NP_D375N_ mutation. Since MxA protein is usually not expressed in laboratory mice, these findings suggest that a virus variant containing the NP_D375N_ mutation would not have a significant disadvantage in replicating in Mx-positive human cells unlike those variants harbouring the NP_G102R_ and NP_M105K_ mutations.

**Figure 3.**
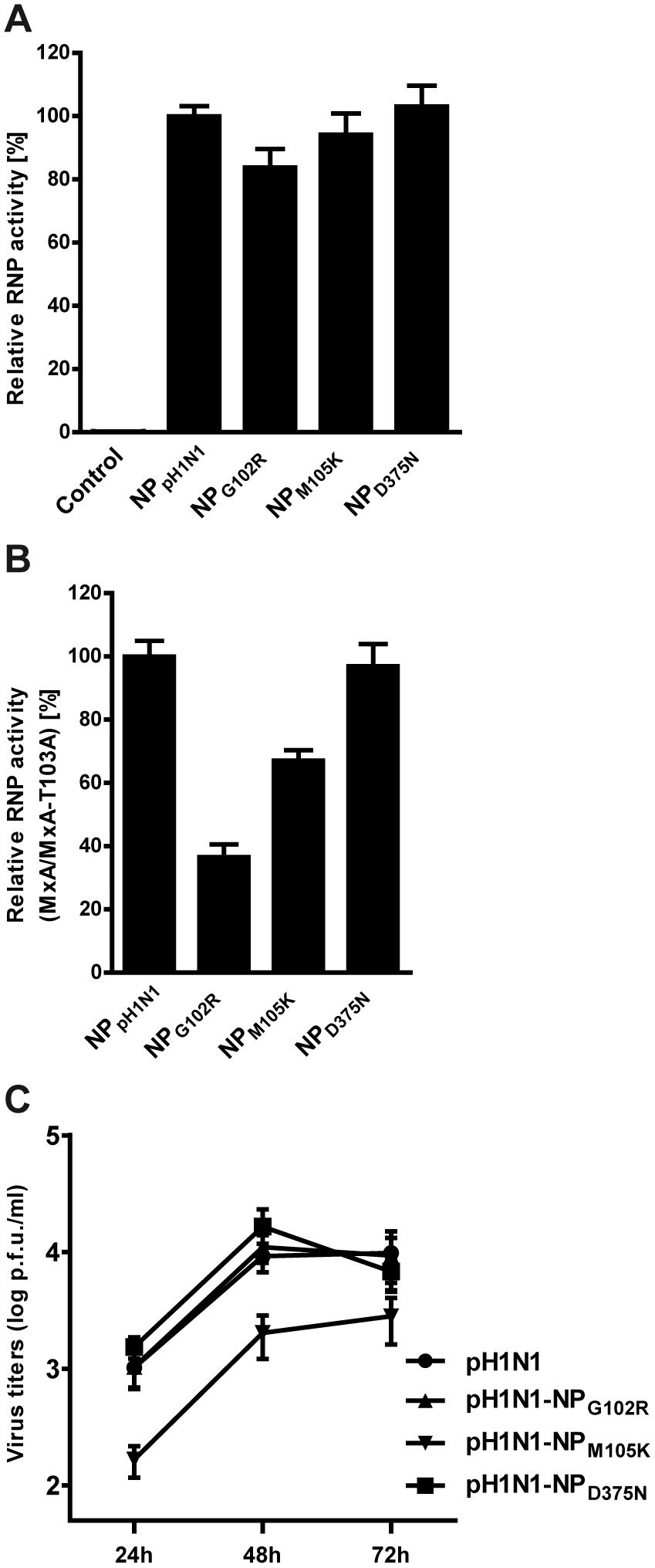
Effect of NP mutations on pH1N1 viral replication. **A.** Biological activity of recombinant pH1N1 RNPs with adaptive NP mutations. Human HEK cells were co-transfected with plasmids encoding for PB1, PA, PB2 and NP of the pH1N1 virus, as well as with reporter constructs encoding for the *Renilla* luciferase under a constitutive promoter and for the firefly luciferase gene in negative polarity, flanked by the non-translated regions of the influenza NP segment. As a negative control plasmid expressing PB1 was omitted in the transfection. Plasmids encoding parental or mutant NP were used as indicated. At 20 h post transfection, luciferase activity was determined. Values for firefly luciferase activity were normalized using the *Renilla* luciferase activity and the activity of the pH1N1 RNP was set 100 % (means ± SEM; *n* = 15 biological replicates). According to one-way ANOVA, there was no significant difference between the mutant and the parental pH1N1 RNPs. **B**. Biological activity of recombinant RNPs with adaptive NP mutations in the presence of MxA. Human HEK cells were cotransfected as described in (A) but co-transfection of MxA expression plasmid was included. RNP activity in the presence of the antivirally inactive MxA-T103A mutant was used to normalize the data obtained with MxA. Values were normalized to the activity of the parental pH1N1 RNP (means ± SEM; *n* = 6-9). According to one-way ANOVA, there was a significant difference between the mutant RNPs and the parental one. **C**. Viral replication kinetics in the human epithelial A549 cell line infected with recombinant viruses containing the different NP adaptive mutations. Cells were infected at an MOI of 0.1 with parental (pH1N1) or adapted mutant viruses. Virus titers of the supernatants taken at the indicated hours post infection (p.i.) were determined by plaque assay (means ± SEM; *n* = 7-9 biological replicates). According to one-way ANOVA, there was no significant difference between the mutant and the parental pH1N1 recombinant viruses.

We next analyzed the effect of these NP host adaptive mutations on the virus replication kinetics in human epithelial cells of lung origin (**Fig. 3C**). No significant differences were observed upon infection with recombinant viruses possessing NP_G102R_ or NP_D375N_ mutations compared to the parental WT strain. However, viral replication was impaired in pH1N1-NP_M105K_ infected cells particularly at early time points.

These findings suggest that viral replication kinetics is impaired during early time points of infection with the pH1N1-NP_M105K_ virus mutant.

### Viral escape mutations in NP present increased importin-α7 binding affinity

Next, we addressed the question whether the observed NP adaptive mutations might alter the binding affinity to importin-α isoforms (**Fig. 4A**). We tested NP binding to importin-α1, -α3 and -α7 isoforms which have been shown to play a crucial role in influenza virus replication in mammalian cells and regulating antiviral immune responses depending on virus kinetics [13, 15]. Moreover, these isoforms are considered representative of each of the three subfamilies of mammalian importin-α proteins [19]. Although the NP mutations had no significant effect on the association of NP to any of the importin-α isoforms, the binding of NP to importin-α7 was in general 10-fold higher than to importin-α1 (**Fig. 4B)**. The potential interaction interfaces between NP and importin-α proteins were also analyzed *in silico* (**Fig. 4C**) using a normalized score (i.e. z-score, see methods). The distribution of the z-scores of all the putative interaction configurations correlates with the stability of the contact and thus with the theoretical affinity between these proteins [20]{Marin-Lopez, 2017 #3136}. Differences between the individual importin-α isoforms are difficult to evaluate due to the missing NLS1-motif in the used NP structures. Regardless, when comparing the individual mutants in each isoform, the distributions of z-scores for the interaction between importin-α1, -α3 or -α7 and the parental NP aligned with the biological findings regarding the interaction of importin-α isoforms with the mutant NP variants.

**Figure 4.**
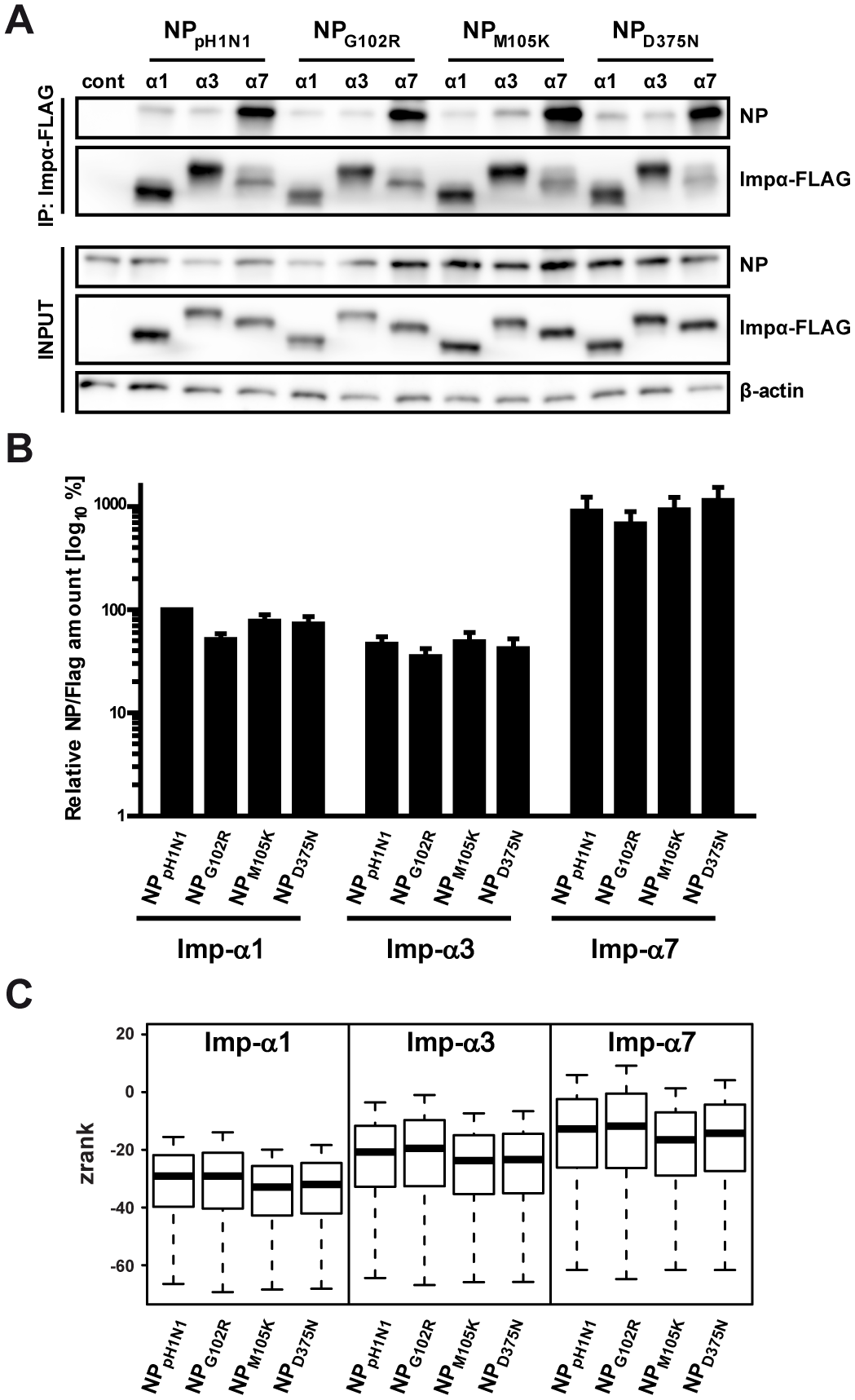
Importin-α binding affinities to NP monomers harboring α7^−/−^ adaptive mutations. **A**. Human HEK cells were co-transfected with plasmids encoding for FLAG-tagged human importin-α isoforms (α1, α3 or α7) and with plasmids for parental or mutant pH1N1 NP as indicated. NP-only transfected cells served as a control (cont). At 48 h post transfection, importins were immunoprecipitated using the FLAG-tag and the amount of coimmunoprecipitated NP was determined by Western blot analysis. β-actin was used as a loading control in the input samples. **B**. Quantification of NP binding to over-expressed FLAG-tagged importin-α isoforms. Values for NP amounts were normalized to the precipitated importin-α amount and the relative amount of NP bound to importinα1 was set 100 % (means ± SEM; *n* = 3 biological replicates with 3-4 technical replicates each). When comparing the affinity for each importin-α isoform, there was no significant difference between the mutant and the parental pH1N1 NP according to one-way ANOVA. **C**. Docking analysis of the interaction of the NP variants with the different human importin-α isoforms. The box-and-whisker plots represent the Zrank score distribution of the top scored 1000 decoys for each NP-importin-α combination. Lower Zrank values represent decoys with a higher probability to mimic the native interacting conformation.

Thus, although individual viral escape mutations in NP do not affect importin-α binding affinities under these conditions, all showed higher interaction affinity to importin-α7.

### Viral escape mutations in NP alter importin-α requirement during infection

Next, we analyzed whether these viral escape variants may have led to a dependency on importin-α isoforms other than importin-α7 [16]. Therefore, we studied the replication of recombinant NP viral escape mutant viruses in importin-α^−/−^ murine embryo fibroblasts (MEFs) isolated from WT, α1^−/−^, α3^−/−^ or α7^−/−^ mice, which were infected with the respective virus variants using single-cycle conditions (**Fig. 5A-D**). Consistent with previous reports [16], we could confirm that the dependency on importin-α7 remained unaltered even upon adaptation to α7^−/−^ mice since virus replication in the absence of importin-α7 was always lower than in WT cells. The requirement for other importin-α isoforms was different depending on the NP adaptive mutation. While replication efficiency of the parental pH1N1 virus in α3^−/−^ or WT cells was similar, pH1N1-NP_G102R_ and pH1N1-NP_D375N_ viruses replicated less efficiently when importin-α3 was not expressed. In contrast, the NP_M105K_ mutation increased virus replication in the absence of importin-α3 and also in the absence of importin-α1. Interestingly, importin-α1 expression did not alter the replication properties of the pH1N1-NP_G102R_ and pH1N1-NP_D375N_ viruses.

**Figure 5.**
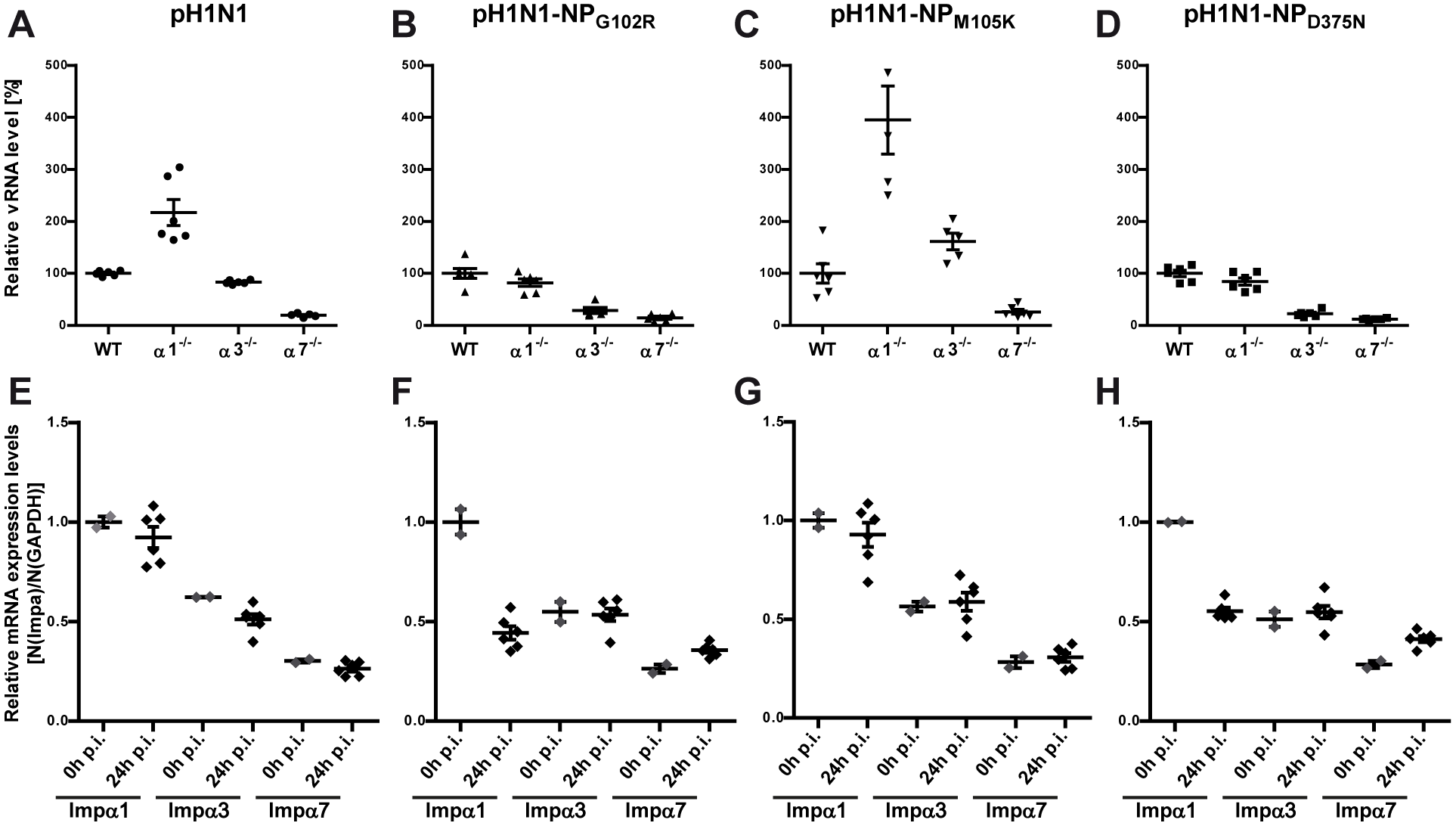
Effect of NP mutations on importin-α requirement during pH1N1 virus replication. **A-D**. Viral RNA accumulation during infection in vitro. WT, importin-α1^−/−^, -α3^−/−^ and -α7^−/−^ MEFs were infected at an MOI of 1.5 with parental (pH1N1 in A) or adapted mutant viruses (pH1N1-NP_G102R_ in B, pH1N1-NP_M105K_ in C, pH1N1-NP_D375N_ in D). After 24 h, total RNA was isolated and the amount of NP vRNA determined by RT-qPCR. Total RNA amount was used for normalization. vRNA accumulation upon infection of the different importin-α^−/−^ MEFs is shown relative to the NP vRNA levels in WT MEF which was set 100 % for each virus tested. Each data point represents an individual sample (means ± SEM; *n* = 5-6 biological replicates). **E-H**. Importin-α mRNA expression levels (α1, α3, α7) in WT MEFs infected with the parental (pH1N1 in E) or adapted mutant pH1N1 viruses (pH1N1-NP_G102R_ in F, pH1N1-NP_M105K_ in G, pH1N1-NP_D375N_ in H). RT-qPCR data were obtained using mouse specific importin-α primers. Relative expression values of importin-α1 were set 1 after normalization against murine GAPDH as a reference. Each data point represents an individual sample (means ± SEM; *n* = 2-6 biological replicates).

In order to assess whether these effects on viral RNA maybe affected by altered importin-α transcription, we measured importin-α mRNA levels in infected cells (**Fig. 5E-H**). Herein, particularly infection with the pH1N1-NP_G102R_ and pH1N1-NP_D375N_ viruses mediated reduced importin-α1 mRNA levels by 2-fold (**Fig. 5F, H**). These findings correlate with unaltered replicative fitness of pH1N1-NP_G102R_ and pH1N1-NP_D375N_ viruses in the presence or absence of importin-α1 (**Fig. 5B, 5D**).

Thus, these results suggest that NP_G102R_ and NP_D375N_ adaptive mutations mediate increased virus dependency on importin-α3 while importin-α1 mRNA levels increase. However, the NP_M105K_ adaptive mutation is particularly affected by the lack of importin-α1 and -α3 expression.

### Interface Frequency Profiles between NP and importin-α proteins are different depending on the isoforms and the host species

Next, we assessed NP residues that may be directly involved in the interaction with importin-α proteins. Therefore, we applied *in silico* binding affinity analysis for NP and its adaptive variants against all members of the importin family (**Fig. 6**). For human and murine species, similar IFPs (Interface Frequency Profiles) were observed for NP interaction with importin-α1 and -α8; less with importin-α3 and -α4 and less, but still similar, with importin-α5 and -α7. The IFP of NP with importin-α5 and -α7 is more scattered than with other isoforms. This suggests that the interaction between NP and importin-α5 or -α7 will require a longer period of exploration, lowering the velocity of binding.

**Figure 6.**
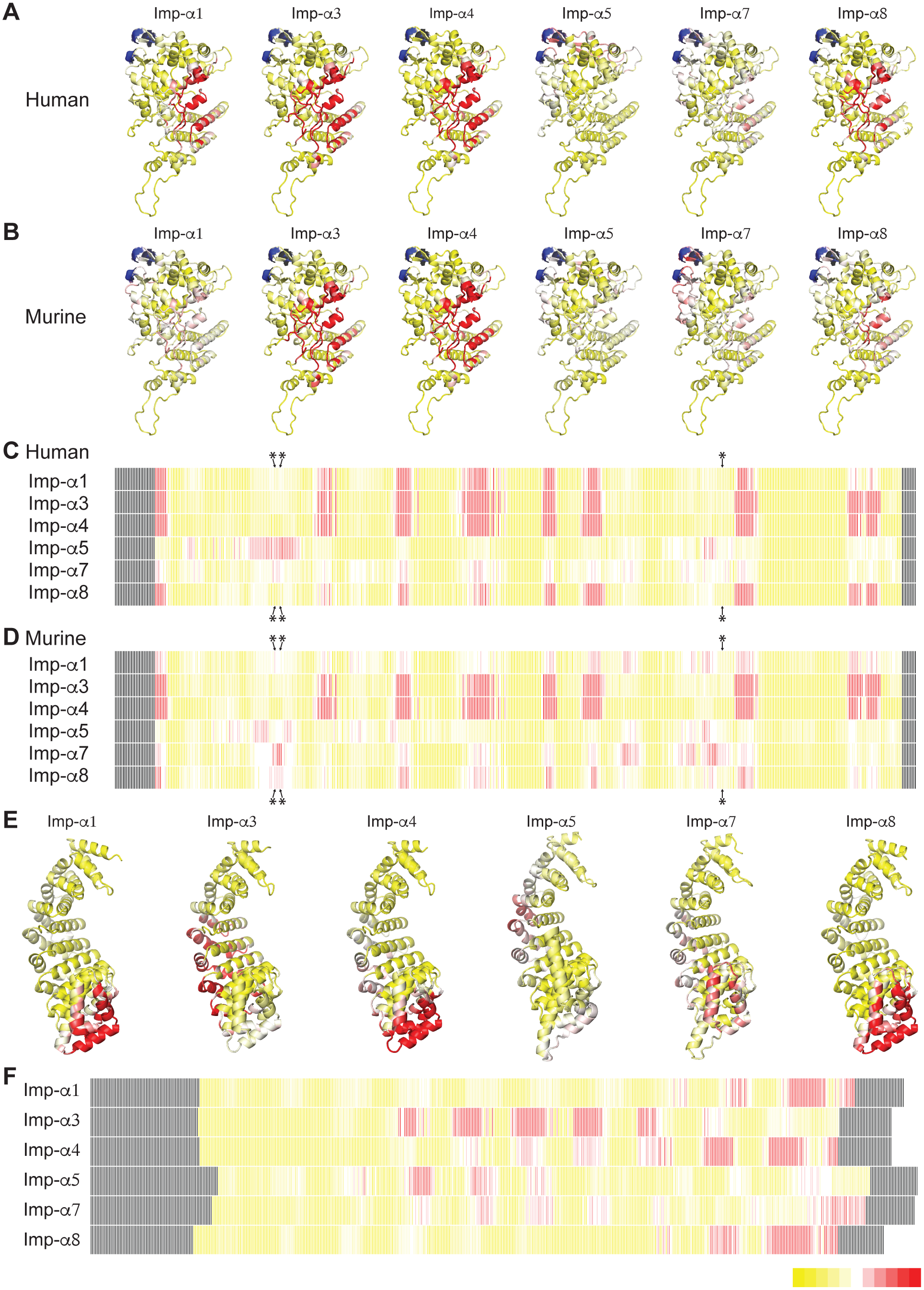
Putative interaction interfaces of NP with the different importin-α isoforms. Interface frequency profiles (IFP) depicted through residue colour as shown in the bottom right legend (yellow represents less and red more frequently found positions in the interaction interface). Each cell of the matrix represents a residue. Matrix cells in black represent the residues that were not present in structure used for the docking analysis. Positions of the adaptive mutation sites (G102R, M105K and D375N) are highlighted in blue in parental pH1N1 NP structure or with asterisks on NP sequence. **A.** IFPs over the structural model of NP with the different human importin-α isoforms. **B.** IFPs over the structural model of NP with the different murine importin-α isoforms. **C.** IFPs over the sequence of NP with human importin-α isoforms. **D.** IFPs over the sequence of NP with murine importin-α isoforms. **E.** IFPs over the structural models of each human importin-α isoform. **F.** IFPs over the sequence of each human importin-α isoform.

This analysis showed that the different IFPs of pH1N1-NP (a) mirror the classification of importin subfamilies [21] and (b) that there is a higher divergence of the α1 subfamily (-α1, -α8) between species than with any of the others. The analysis of IFPs of importin-α isoforms shows different potential binding regions (**Fig. 6E-F**). In all the cases, the region of the arm repeat that forms the concave surface of the importin shows no actual contacts with any decoy because this region has been identified as the binding site for NLS domains [22] that lacks in our NP model. Importin-α4 and -α7 show a scattered pattern of low-frequency residues across almost all the arm repeats. Importin-α3 shows a discrete pattern of high-frequency residues across a set of consecutive arm repeats. Importin-α5 presents a discrete pattern englobing arm repeats from 3^rd^ to 5^th^. Importin-α1, -α4 and -α8 present distinct discrete pattern in the 10^th^ arm repeat.

These high frequency discrete patterns indicate that the importin arms are most likely performing a secondary non-NLS mediated interaction with NP.

### NP viral escape mutations alter the putative interaction interface of NP with the different importin-α isoforms

Then, we selected one importin-α isoform for each subfamily to study the effect of NP viral escape mutations on the interaction frequency profiles (IFP) with human and murine importin-α isoforms (**Fig. 7**). We observed that IFPs of NP for the interactions with importin-α1 and -α7 isoforms became more discrete, with slightly different profiles for all mutations, while for importin-α3, the effect was negligible and showed similar profiles for all mutations. Changes on the IFP depended on the NP mutation, but also on the species and importin-α isoform, and were especially relevant for the NP_G102R_ variant. When this mutation was introduced, the IFPs of NP for the interactions with importin-α1 and importin-α7 displayed higher frequency around the mutation sites (positions 102, 105 and 375), but the profile of the interaction with importin-α3 was mostly unaffected. The effect of NP_M105K_ was minor, while interestingly NP_D375N_ mutation increased the scattered behaviour of the IFP for importins -α1, -α3 and -α7 and for both human and murine species.

**Figure 7.**
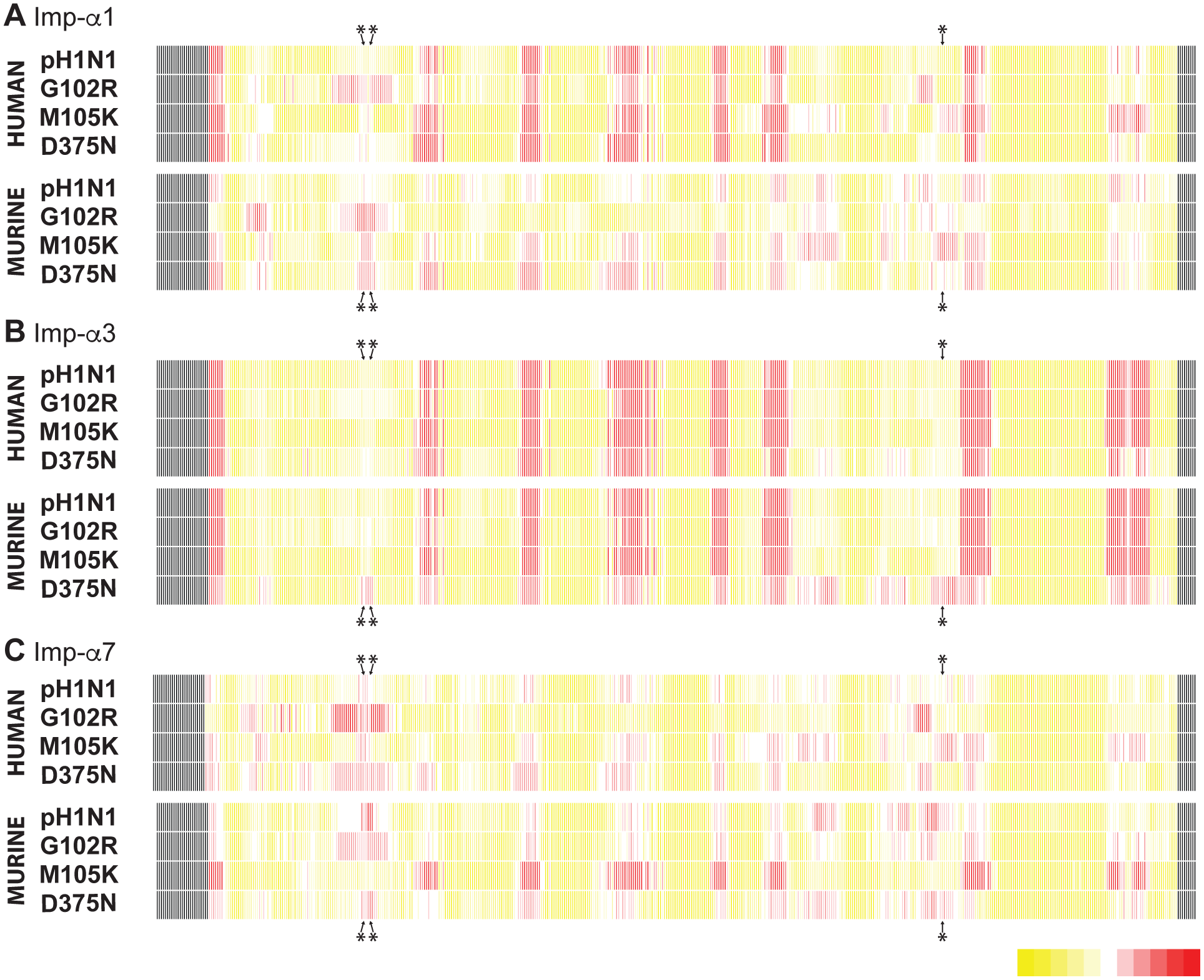
Effect of NP adaptive mutations on the putative interaction interfaces of NP with the different importin-α isoforms. Interface frequency profiles (IFP) mapped over the sequence of parental pH1N1 NP and those of the adaptive mutant variants (G102R, M105K and D375N). Each cell of the matrix represents a residue. Matrix cells in black represent the residues that were not represented in the NP structure used for the docking analysis. Frequency is depicted by residue colour as shown in the bottom right legend (yellow represents less and red more frequently found positions in the interaction interface). Positions of the adaptive mutation sites (G102R, M105K and D375N) are highlighted with asterisks. **A.** IFPs of parental or mutant NP with human and murine importin-α1 isoforms. **B.** IFPs of parental or mutant NP with human and murine importin-α3 isoforms. **C.** IFPs of parental or mutant NP with human and murine importin-α7 isoforms.

These findings suggest that adaptive mutations in NP are likely to alter putative interaction sites on the protein surface which may then mediate nuclear import by the classical transport machinery.

## DISCUSSION

Since influenza virus replication and transcription occurs in the nucleus of the infected cell, the nuclear import machinery and the importin-α proteins in particular, play a key role during infection [3]. In order to better understand the role of importin-α7 during mammalian adaptation of influenza viruses, we investigated 2009 pandemic H1N1 virus (pH1N1) adaptation in a restrictive environment where the most relevant isoform of these host factors was not expressed [16]. Although many polymorphisms were found at low frequency in several viral genes during this adaptation process, only three independent NP variants were found at high frequency and already at early stages of the process.

Virus evolution occurs under specific selective pressures and can provide an advantage for the virus in that specific context and environment. Unexpectedly, dependency on importin-α7 expression still remained upon adaptation of pH1N1 to importin-α7^−/−^ mice, highlighting the importance of this host factor for influenza virus replication. This phenomenon was also observed in our previous study, where we characterized a virus that became highly pathogenic for α7^−/−^ mice by acquiring mutations in the viral RNP as well as in the surface glycoproteins HA and NA [16]. Additionally, the C57BL/6J strain used in this study does not, similar to many laboratory inbred mouse strains, express a functional Mx1 protein, the murine homologue of human MxA (IFN)-inducible nuclear protein [23]. Therefore, the virus adaptation performed here occurred in the absence of this antiviral factor. This might allow the occurrence of mutations that in the end could be disadvantageous for the virus in the presence of MxA, as observed in our present study. Furthermore, this suggests a potential relationship between the effects of importin-α7 and MxA on NP during influenza virus infection.

Strikingly, the three adaptive mutations found in NP are located outside of the regions previously described as NLSs and do not affect its structural conformation [16]. Similarly, another mutation in the NP_M105_ residue has been described to provide a new transport-competent nuclear localization signal [24]. Interestingly, our results indicated that these escape mutations occurred independently and triggered different viral replication phenotypes in the absence of other importin-α isoforms from the same nuclear import pathway. Therefore, they could provide different escape mechanisms including secondary interacting interfaces between NP and importin-α arm repeat domains [25, 26].

Hereby, we have been able to assess the variable affinity of the different mutants to several members of the importin-α family with a docking approach through two different strategies [20, 27]. Initially, we evaluated the differences in affinity between each NP mutant and the different importin-α isoforms. Firstly, we observed that the affinity profile of each NP mutant in the computational experiment mimics the experimental evidence, with no significant differences amongst the mutants. Secondly, none from these top decoys showed the concave region of importin-α (formed by the Arm repeats) as a putative binding interface on the side of the importins. This is especially relevant as this region has been described as the NLS1 recognition site. Moreover, we plotted the frequency in which each residue of NP appears in the putative interface for the top 50 scored decoys. We named this plot Interface Frequency Profile (IFP) and interpreted that changes in the profile shape would correspond to changes in the kinetics of the interaction (see methods). Thus, a more scattered IFP in the three-dimensional space requires a longer time to find its final interaction conformation. As a matter of fact, this kind of effect can be observed for the pH1N1-NP_D375N_ mutant, especially at its interaction with importin-α1 and -α7, phenomenon that could clearly affect the efficiency of NP-importin interaction. Importin-α7 showed a more scattered IFP in NP, implying longer kinetics in the interaction of NP with importin-α7, as suggested before for PB2 and importin-α proteins [19]. Parallel results were obtained for importin-α5, as expected due to the sequence similarity to importinα7. Essentially, there are two potential interaction regions of NP with importin-α7: one region coinciding with the potential interaction interface with importin-α3 and importin-α4 and a second region near the locus where the escape mutations are located. Interestingly, the region of this locus is an inhibitory target for MxA while the other three-dimensional section detected is placed just over and after the second segment of the NLS2, indicating that this might be part of a bigger recognition site.

The influenza replication cycle is a complex process where many host factors, including the importin-α proteins [3], play one or several important roles during infection. The here investigated NP_G102R_, NP_M105K_ and NP_D375N_ adaptive mutations alter the properties of this viral protein in different ways and are associated with increased pathogenicity upon adaptation to mice lacking the importin-α7 gene. The generated IFPs differ especially between the NP_M105K_ variant and the other variants. NP_M105K_ showed a profile similar to that of the parental NP, except for higher frequencies when interacting with murine importin-α1 and even higher frequencies with murine importin-α7 when compared to the respective human isoforms. This effect correlates with higher virus replication in murine cells in the absence of importin-α1 and -α3 which would serve as restriction factors for these virus variants. On the other hand, the change in the potential interaction profiles of NP_D375N_, and especially NP_G102R_, with importin-α1 correlates with the suppression of the negative effect that the importin-α1 protein exerts on pH1N1 virus replication in murine cells. This finding correlates moreover also with the observed lower mRNA expression levels of importin-α1 upon pH1N1-NP_D375N_ and -NP_G102R_ infection, suggesting diverse possibilities for the virus to circumvent the absence of importin-α7.

In summary, in this study, we described new regions in the viral NP protein involved in mediating interaction with the importin-α proteins. Diverse elements of the viral NP protein as well as of the host’s importin-α proteins contribute to their interaction and therefore to the functionality of the protein complex, although they are not specifically located at the interaction site.

## MATERIALS AND METHODS

### Cells

Madin-Darby canine kidney (MDCK) cells were grown in minimal essential medium (PAA, Austria) supplemented with fetal calf serum (FCS), L-glutamine, and penicillin-streptomycin. Human embryonic kidney 293T (HEK) and human alveolar adenocarcinoma (A549) cells were grown in DMEM (Dulbecco’s modified Eagle’s medium) supplemented with 10 % FCS, 1 % L-glutamine and 1 % penicillin-streptomycin. WT, α1^−/−^, α3^−/−^ and α7^−/−^ murine embryonic fibroblasts (MEF) were grown in DMEM supplemented with 10 % FCS, 1 % penicillin/streptomycin, 1 % L-Glutamine, 1 % nonessential amino acids (PAA, Austria) and 1 % sodium pyruvate. They were prepared from murine embryos harvested from pregnant females and cells were immortalized by repeated passaging. MEFs were kindly provided by M. Bader (Max Delbrück Center for Molecular Medicine, Berlin, Germany) [28]. All tissue culture reagents were supplied by PAA, Austria.

### Adaptation of pH1N1 influenza virus to importin-α7^−/−^ mice and deep sequencing of viral RNA

The 2009 pandemic H1N1 A/Hamburg/NY1580/09 virus strain (abbreviated as pH1N1) was passaged five times in importin-α7 (α7^−/−^) mice in our previous study [16]. Briefly, we infected three α7^−/−^ mice intranasally with 10^5^ p.f.u. of pH1N1 virus. Three days later, lungs of these mice were excised and the lung homogenates pooled and used to infect a second group of three α7^−/−^ mice. Note that no new mouse experiments were conducted for this study. In this study, virus populations in the lung homogenates as well as the parental virus strain were amplified in MDCK cells followed by virus concentration using the PEG Virus Precipitation Kit (BioVision, USA). Total RNA was extracted with TRIzol reagent (Invitrogen, USA) according to the manufacturer’s instructions.

For each virus sample, 1 µg of total RNA was Dnase I (Life Technologies) treated and purified using RNeasy MinElute columns (Qiagen, Netherlands). Sequencing libraries were generated with the ScriptSeq v2 RNA Seq Kit (Epicentre, USA) using 150 ng of purified RNA as input as recommended by the manufacturer. Library fragments between 200 and 600 nt were size-selected using Pippin Prep (Sage Science, USA) and further purified with Agencourt AMPure XP beads (Beckman Coulter, USA). Size and quality of the libraries were visualized on a BioAnalyzer High Sensitivity DNA Chip (incl. in High Sensitivity DNA Kit, Agilent Technologies, USA). Diluted libraries (2 nM) were multiplex-sequenced on the MiSeq platform (Illumina, USA) generating between 2M and 4M paired-end reads.

Reads were aligned to the reference assembly of A/Hamburg/NY1580/09 (H1N1) (GenBank: HQ104928.1, HQ104926.1, HQ104924.1, HM598305.1, HQ104929.1, HQ104927.1, HQ104925.1, GU480807.1) using the Burrows Wheeler Aligner (v0.7.5a) [29]. SAMtools (v0.1.19) [30] was employed for format conversions and the removal of putative PCR duplicates. Variant calling was performed with SAMtools and VarScan2 (v2.3.6) [31], not considering bases with Phred scores lower than 30. Finally, called variants were annotated with SnpEff (v3.3) [32]. Sequence data reported in this publication have been submitted to the European Nucleotide Archive (ENA) and are available at http://www.ebi.ac.uk/ena/data/view/PRJEB8023.

### Plasmids and recombinant pH1N1 viruses

The pHW2000 rescue plasmids and pcDNA3.1 expression plasmids [16] encoding for the nucleoprotein (NP) of pH1N1 were used to introduce the specific adaptive mutations by site-directed mutagenesis. Then, these mutated pHW2000 vectors were used to generate the recombinant influenza viruses pH1N1-NP_G102R_, pH1N1-NP_M105K_ and pH1N1-NP_D375N_ by reverse genetics as described before [33]. Virus stocks were propagated in MDCK cells and the sequence of the encoded NP gene was verified.

The pCAGGS-MxA and pCAGGS-MxA-T103A plasmids were kindly provided by Martin Schwemmle (University of Freiburg, Germany) [18]. The pcDNA-importin-α1-FLAG, pcDNA-importin-α3-FLAG and pcDNAimportin-α7-FLAG constructs were described previously [14].

### Polymerase activity

HEK cells seeded in 12-well plates were co-transfected with 10 ng of pCDNA3.1 plasmids encoding for PB1, PB2 and PA plus 100 ng of NP-encoding plasmid (parental NP or indicated mutant) to generate recombinant RNPs. To standardize the transfection experiments, constructs pPol-I-NP-Luc-human (100 ng; encoding the firefly luciferase in negative polarity flanked by the non-translated regions of the influenza NP segment) [34] and pRL-TK (30 ng; encoding the *Renilla* luciferase; Promega, USA) were co-transfected as well. To evaluate the effect of the NP adaptive mutations on the resistance to antiviral MxA, 200 ng of MxA- or MxA-T103A-encoding plasmids were co-transfected as described before [18]. An empty pCDNA3.1 vector construct was added to achieve equal amounts of transfected DNA per well. The transfection solution was prepared by incubating the DNA mixture with polyethylenimine (PEI; Polysciences) and DMEM in a ratio of 2.4 μg PEI to 600 μl DMEM for 20 min at room temperature. At 20 h post transfection, luciferase activity was measured using a Dual-Luciferase Reporter Assay System (Promega, USA) according to the manufacturer’s instructions. Successful MxA expression was verified by Western blot (data not shown). All experiments were performed in biological triplicates.

### Virus growth kinetics

A549 cells were infected at an MOI of 0.1 with pH1N1, pH1N1-NP_G102R_, pH1N1-NP_M105K_ or pH1N1-NP_D375N_ viruses in the presence of 0.25 μg/μl L-1-tosylamide-2-phenylethyl chloromethyl ketone (TPCK)-trypsin. Supernatants were collected at 0, 24, 48 and 72 h post infection (p.i.) and virus titers were determined as plaque forming units per ml (p.f.u./ml) on MDCK cells.

### Co-immunoprecipitation assay

HEK cells seeded in 12-well plates were co-transfected with pCDNA3.1 expression vectors encoding for the viral NP protein (3 μg of plasmid DNA) and each of the importin-α isoforms (700 ng of plasmid DNA). After 48 h, cells were lysed by incubation in 300 μl of buffer L1 (50 mM Tris-HCl, 100 mM NaCl, 5 mM EDTA, 0.5 % Igepal, 1 mM dithiothreitol [DTT], 1x HALT™ Protease and Phosphatase Inhibitor Cocktail, 1x EDTA Solution (both: 100x, Pierce/Thermo Scientific, USA) for 2 h at 0 °C with occasional vortexing. Then, the lysate was centrifuged for 30 min at 12,000 rpm and 4 °C. Immunoprecipitations were performed using the EZview Red ANTI-FLAG M2 affinity gel (Sigma, USA) and eluted by adding 3x FLAG peptide (Sigma, USA) according to the manufacturer’s instructions. Quantification of co-immunoprecipitation products was performed by Western blot analysis using mouse anti-FLAG (cat. no. F3165, Sigma, USA) and rabbit anti-FPV serum [35] antibodies. The β-actin antibody (Abcam, UK) was used for normalization of the total protein amount used in respective cell lysates. Immunoreactive bands were visualized with the Bioimager Image Quant LAS 4000 at non-saturated levels and quantified by densitometry with ImageJ software.

### mRNA expression analysis in infected MEFs

Confluent monolayers of WT, α1^−/−^, α3^−/−^ and α7^−/−^ MEFs were infected at an MOI of 1.5 with pH1N1 recombinant viruses containing NP mutations as indicated for 1 h at 37 °C. The virus inoculum was aspirated and growth media, without FCS but supplemented with 0.2 % bovine serum albumin, was added. Total RNA was isolated at 24 h p.i. using the TRIzol reagent (Invitrogen, USA) according to the manufacturer’s instructions.

Viral RNA amount was quantified by real-time quantitative PCR (RT-qPCR). Therefore, viral RNAs were reverse transcribed using primers specific for NP vRNA (5’-ccgacatgcgaacagaagt-3’) with SuperScript III reverse transcriptase (Invitrogen) at 50 °C for 1 h. The resultant cDNAs were then quantified by using a set of RT-qPCR primers specific for NP (fw: 5’-ccgacatgcgaacagaagt-3’ and rev: 5’-ccgagagctcgaagactcc-3’), the hydrolysis probe #129 (cat. no. 04693655001, Roche) and the LightCycler 480 Probes Master mix (Roche) in the LightCycler 96 Real-Time PCR system (Roche, Switzerland) with endpoint fluorescence detection: 10 min at 95 °C, 45 amplification cycles (10 seconds at 95 °C and 30 seconds at 60 °C). Reactions were set up in technical triplicate and obtained data was analyzed by using the LightCycler 96 software. The standard curve was obtained by analyzing serial dilutions of linearized pC-HH15-NP plasmid [16]. Total RNA present in cell lysates was used for normalization.

Importin-α mRNA expression levels in infected WT MEFs were also quantified by RT-qPCR. Specifically, cDNA was generated using random nonamer primer (Gene Link™, pd(N)9, 26-4000-06) and the SuperScript™ III Reverse Transcriptase (Invitrogen) according to the manufacturer’s instructions. Then, RT-qPCR specific for importin-α1, -α3, and -α7 genes and the reference gene GAPDH was performed with specific oligos (importin-α1: fw: 5’-GCGTCGCCGCCGAATAGAAGTTAATG-3’ and rev: 5’-TTCCGAGCAGCTTGAGTAGCTTGGAG-3’; importin-α3: fw: 5’-ACACTTCCCAGCACTCCTCACTCATC-3’ and rev: 5’-AAGATAGGCCACTTGGTCTTTCCTTCC-3’; importin-α7: fw: 5’-TGACTTGCAGTTAGCAACCACGCAG-3’ and rev: 5’-ATGGGGACAGCTCCTGCTTCAATG-3’; GAPDH: fw: 5’-CCACTGAAGGGCATCTTGGGCTAC-3’ and rev: 5’-GGTGGGTGGTCCAGGGTTTCTTAC-3’;). Singleplex reactions of 10 µl were set up manually in UltraPure DNase/RNase-Free Distilled Water (Gibco) in MicroAmp^®^ Optical 96-Well Reaction Plates (Invitrogen, #4306737): 5 µl Platinum^®^ SYBR^®^ Green qPCR SuperMix-UDG (2x, Invitrogen), 0.02 µl ROX Reference Dye (25 µM, Invitrogen, final concentration: 50 nM), 300 nM of forward and reverse primer each, and 1 µl cDNA template. RT-qPCR runs were conducted on the ABI 7500 Fast System (Applied Biosystems) in the Standard 7500 mode with endpoint fluorescence detection: 2 min at 50 °C, 3 min at 95 °C, 50 amplification cycles (15 seconds at 95 °C, 10 seconds at 65 °C, and 30 seconds at 72 °C). Analysis was performed in triplicate or quadruplicate for each gene in each sample. A melting curve analysis was performed after the RT-qPCR run on the ABI 7500 Fast System (15 seconds at 95 °C, 1 min at 60 °C, 15 seconds at 95 °C). Additionally, generation of amplicons with the correct size was checked by agarose gel electrophoresis. Data of reactions with false products were excluded from subsequent data analyses. For the calculation of relative expression values, the E^−ΔΔCT^ method was used [36, 37]. The Rn-values of the RT-qPCR were exported from the SDS Software v1.3.1 (Applied Biosystems) to Microsoft Office Excel 2007 and N_0_-values for the starting concentration of the transcript in the original sample were obtained using the LinReg PCR Software v11.185 [37].

### Molecular modeling of NP interaction with importin-α isoforms

The computational study of the interaction between NP or its mutant variants and the different isoforms of the importin-α family was carried out through molecular docking with PatchDock [38]. The structural models for NP and its mutants (NP_G102R_, NP_M105K_ and NP_D375N_) were generated as described in our previous work [16]. The models lack the first 25 residues and the last 9 residues due to the lack of templates for that particular segment. This does not affect the comparison of the effects of each NP mutant over a particular importin isoform. Importin-α isoform with known structures were retrieved from the Protein Data Bank [39]. This was the case for human importin-α1 (KPNA2; 4WV6 [40]), murine importin-α1 (KPNA2; 4UAF [19]), human importin-α3 (KPNA4; 4UAE [19]), human importin-α5 (KPNA1; 2JDQ [41]) and human importin-α7 (KPNA6; 4UAD [19]). Due to the high level of conservation among the members of the importin-α family, the rest of the required structures (human importin-α4 (KPNA3) and -α8 (KPNA7) as well as murine importin-α3, -α4, -α5, α7 and -α8) were obtained by homology-modelling and retrieved from the SWISS-MODEL server [42]. The importin-β-binding (IBB) domain of both known structures and models was excluded from our computational studies as the highly flexible peptide region between the two importin domains impedes a reliable way to model their relative position.

Docking was performed for all members of the importin-α family against all of the different NP variants using PatchDock with the higher accuracy sampling parameters. This produced between 54.000 and 90.000 poses, i.e. putative interacting configurations, for each protein-protein interaction. Hydrogens were added to the models with Reduce [43] and the individual energy of each pose was analyzed with Zrank [44].

Docking techniques have been proven able to identify interfaces between interacting proteins and as a tool to assess the probability of a protein-protein interaction [20]. Despite that, it is difficult to pinpoint the pose that better represents the three-dimensional interaction between two proteins, as current methods tend to score native and near-native conformations amongst the best-scored conformations together with other poses [45]. To overcome this issue, we adapted Li *et al.* approximation [27] by building Interface Frequency Profiles (IFP).

IFPs are constructed by calculating the frequency in which each amino acid of two interacting proteins occur in the interface of the 50 top scored poses obtained from docking. We will differentiate between two types of IFP: discrete and scattered.

In a discrete IFP some patches of amino acids in the surface of a protein have a conspicuously higher frequency than the rest. A scattered IFP is when multiple three-dimensional regions have similar frequencies. In agreement with the funnel-like intermolecular energy landscape theory [46], finding the native pose between two interacting proteins requires a longer and time-consuming search for a protein with scattered IFP than for a protein with discrete IFP. Thus, we hypothesize that if a protein has two different IFPs (which depend on the interacting partners), one scattered and one discrete, the difference is not necessarily shown in the binding energy, but in the time of exploration towards the native pose, and consequently in the interaction kinetics.

### Statistical analysis

All data shown are presented as mean ± SEM. Mean, SEM, and ANOVA were calculated with Prism GraphPad software (GraphPad Software, Inc., USA).

## Acknowledgments

We are grateful to Daniela Indenbirken for excellent technical assistance and Prof Adam Grundhoff for providing the service of his unit. This work was supported by grants from the European Union (FLUPHARM to GG), the Emmy-Noether Programme of the German Research Foundation (GA 1575/1-1 to GG) and the Spanish Ministry of Science and Innovation (FEDER BIO2011-22568, EUI2009-04018 to BO). PRI was funded by the Alexander von Humboldt Foundation (3.3SPA/1142463 STP-2). JB was supported by a MICINN fellowship (FEDER BIO2008-0205). The Heinrich Pette Institute, Leibniz Institute for Experimental Virology, is supported by the Free and Hanseatic City of Hamburg and the Federal Ministry of Health. The funders had no role in study design, data collection and analysis, decision to publish, or preparation of the manuscript.

## Author contribution

PRI and GG designed the study and wrote the paper. PRI and MA performed the experiments shown in Figures 1 and 2. PRI performed the experiments shown in Figures 3 and 4. PRI and ST performed the experiments shown in Figure 5. PRI and JB designed and performed the experiments shown in Figures 6 and 7. All authors analyzed the results and approved the final version of the manuscript.

